# Mapping the organization and dynamics of the posterior medial network during movie watching

**DOI:** 10.1101/2020.10.21.348953

**Authors:** Rose A. Cooper, Kyle A. Kurkela, Simon W. Davis, Maureen Ritchey

## Abstract

Brain regions within a posterior medial network (PMN) are characterized by sensitivity to episodic tasks, and they also demonstrate strong functional connectivity as part of the default network. Despite its cohesive structure, delineating the intranetwork organization and functional diversity of the PMN is crucial for understanding its contributions to multidimensional event cognition. Here, we probed functional connectivity of the PMN during movie watching to identify its pattern of connections and subnetwork functions in a split-sample replication of 136 participants. Consistent with prior findings of default network fractionation, we identified distinct PMN subsystems: a Ventral PM subsystem (retrosplenial cortex, parahippocampal cortex, posterior angular gyrus) and a Dorsal PM subsystem (medial prefrontal cortex, hippocampus, precuneus, posterior cingulate cortex, anterior angular gyrus). Ventral and Dorsal PM subsystems were differentiated by functional connectivity with parahippocampal cortex and precuneus and integrated by retrosplenial cortex and posterior cingulate cortex, respectively. Finally, the distinction between PMN subsystems is functionally relevant: whereas both Dorsal and Ventral PM connectivity tracked the movie content, only Ventral PM connections increased in strength at event transitions and appeared sensitive to episodic memory. Overall, these findings reveal PMN functional pathways and the distinct functional roles of intranetwork subsystems during event cognition.

## 1. Introduction

Complex cognitive processes, such as understanding and remembering events, rely on the functional interactions of brain networks, which are defined as groups of structurally and functionally connected brain regions. One such network is the posterior medial network (PMN), which consists of regions that are strongly functionally connected with the parahippocampal cortex, including posterior medial temporal lobe, medial and lateral posterior parietal cortex, and medial prefrontal cortex (Libby et al., 2012; Wang et al., 2016). As part of the default network (Buckner et al., 2008; Raichle et al., 2001), the PMN seems to play a pivotal role in event perception and memory, with its network-level functional role described as forming situational or contextual models (Ranganath & Ritchey, 2012; Reagh & Ranganath, 2018; Ritchey, Libby, et al., 2015). An important characteristic of events is that they are multidimensional, including their visuo-spatial content, conceptual significance, and attributed thoughts and emotions. While large-scale networks help to paint a broad picture of regions that tend to affiliate during cognitive tasks, distinct components of cognitive processes are likely associated with smaller subnetworks of brain regions (Cabeza & Moscovitch, 2013), refined from the large-scale network in which they are embedded. Therefore, understanding how the PMN supports the representation of multidimensional events requires a better understanding of its subnetwork architecture (Ritchey & Cooper, 2020). Here, we deconstruct the organization of the PMN and test how its constituent connections relate to event cognition.

Regions of the PMN tend to function in a cohesive manner, exhibiting strong task-independent correlations in BOLD activity within the default network as well as task-related coactivation. Prior research has shown increased activity across the PMN during the recollection and construction of specific events (Benoit & Schacter, 2015; Rugg & Vilberg, 2013; Schacter et al., 2007; Spreng et al., 2009), network-wide multivariate representation of event-specific information (Chen et al., 2017; Robin et al., 2018), as well as reliable PMN responses to transitions between event contexts (Baldassano et al., 2017; Ben-Yakov & Henson, 2018; Reagh et al., 2020). Research that has directly modulated the PMN also supports its cohesive structure: non-invasive brain stimulation of left angular gyrus (AG) — the PMN network region most accessible for stimulation —increases BOLD activity (Kim et al., 2018), and functional connectivity throughout the PMN during episodic tasks (Warren et al., 2019), confirming the strong functional dependence between these regions. Functional communication within the PMN not only increases during event processing, but also dynamically tracks the amount of information later recalled (Cooper & Ritchey, 2019; Simony et al., 2016). The overarching role of PMN communication appears tied to the construction of meaningful contextual frameworks, as evidenced by increasing connectivity among PMN regions as the temporal structure of naturalistic events is learned (Aly et al., 2018). Beyond this research, it is important to highlight the functionally diverse organizational structure of the PMN, a flexibility which may explain its ability to dynamically adapt to varying task demands.

Two emerging lines of evidence suggest that although episodic construction may describe PMN function at the network level, there is also significant diversity of cognitive processes and functional connectivity profiles associated with PMN regions (Ritchey & Cooper, 2020). The first line of research illustrating PMN diversity comes from the multidimensional analysis of mnemonic content, drawn from both standard episodic tasks as well as movie perception and recall. Separable attributes of episodic memory are associated with distinct PMN regions: Whereas medial temporal lobe (MTL) regions facilitate successful retrieval of event information, parietal cortex tracks the richness of that information, with memory imageability and precision dissociating medial and lateral parietal regions, respectively (Richter et al., 2016). Relatedly, the PMN is fractionated by transient activation within the hippocampus and retrosplenial cortex when accessing episodic information, and sustained activation in dorsal medial and lateral parietal cortex during elaboration (Daselaar et al., 2008; Thakral et al., 2017; Vilberg & Rugg, 2012). Interestingly, PMN regions also show variable temporal resolutions of event context signals during movie watching (Baldassano et al., 2017; Keidel et al., 2017). Ventral medial parietal cortex and parahippocampal cortex separate events at short time-scales whereas dorsal medial and lateral parietal cortex separate events at longer time-scales (Baldassano et al., 2017; Chen et al., 2016). Additionally, parahippocampal and ventral parietal signals are stronger when there is a new narrative context, but lateral parietal activity is increased when an existing context is maintained (Keidel et al., 2017). Taken together, such dissociations point to a hierarchical structure of event cognition within the PMN, with specific event information being conveyed from the MTL and ventral parietal cortex to update representations in dorsal and lateral parietal regions.

The second line of research suggesting a diverse PMN organizational structure comes from resting-state analyses that have shown fractionation of the large-scale default network into distinct subsystems. Such research has demonstrated the presence of a cortical MTL network, including parahippocampal cortex and ventromedial parietal cortex, and a more dorsal network including posterior cingulate cortex, prefrontal cortex, and lateral temporal cortex (Andrews-Hanna et al., 2010; Barnett et al., 2020; Braga & Buckner, 2017; Gordon et al., 2020; Kaboodvand et al., 2018). Moreover, activity of these subsystems appears to correlate with distinct, yet related, cognitive domains: An MTL network may be driven by spatial-contextual processes (Baldassano et al., 2016; Silson et al., 2019) and a Dorsal Medial network shows sensitivity to conceptual information and mental states (Andrews-Hanna et al., 2010; Barnett et al., 2020; DiNicola et al., 2020). The PMN identified in studies of event perception and memory includes brain regions that bridge these previously defined default subsystems. Yet, a focused analysis of intranetwork PMN connectivity, where network definition is limited to areas specifically associated with episodic processing, is lacking. Understanding the organization of the PMN could help to shed light on why the aforementioned functional dissociations occur. Specifically, what are the paths of information flow between brain regions that are associated with episodic processing And are there functionally distinct subsystems within the PMN that differentially contribute to event cognition within the PMN, with specific event information being conveyed from the MTL and ventral parietal cortex to update representations in dorsal and lateral parietal regions.

To address these outstanding questions, we analyzed a subset of the Cambridge Centre for Ageing and Neuroscience (CamCAN) dataset (Shafto et al., 2014; Taylor et al., 2017), a large population-representative sample of individuals who underwent a rich behavioral and neuroimaging testing protocol, generating data ideal for the estimation of dynamic changes in functional connectivity of the PMN during movie watching. First, we aimed to identify separable PMN subsystems based on whole-brain voxel connectivity patterns, additionally testing intranetwork PMN functional connectivity to validate the subsystems and to probe region-specific contributions. Second, we tested the functional significance of intranetwork PMN connectivity dynamics in terms of their sensitivity to movie content, including event transitions, and their relation to individual differences in episodic memory.

## 2. Material and Methods

### 2.1. Data

The data analyzed here were obtained from the CamCAN Stage II data repository (Shafto et al., 2014; Taylor et al., 2017): https://camcan-archive.mrc-cbu.cam.ac.uk/dataaccess/. From this dataset, we selected healthy young adult subjects aged 18-40 who are right-handed, native English speaking, and who had completed the movie watching fMRI scan. A total of 154 (80 female, 74 male; mean age = 30.92, SD = 5.64) subjects met this criteria. After data quality checks, detailed in section 2.4, 18 subjects were removed from the sample, leaving 136 subjects for all analyses. Due to this large sample size, we randomly divided subjects into two equal groups (68 subjects per group), equating for age (group 1: mean = 31.06, SD = 5.75; group 2: mean = 31.12, SD = 5.52) and gender (35 females and 33 males per group). All statistical analyses were run first on group 1 only, allowing us to explicitly test the replicability of our results with group 2.

### 2.2. Task

In the MRI scanner, participants watched a 8 minute movie (Shafto et al., 2014). The movie was a shortened episode of Alfred Hitchcock’s “Bang! You’re Dead” (Hasson et al., 2008, 2010) that was cut in a way that retained the central plot (see Ben-Yakov & Henson, 2018). To allow us to quantify meaningful changes in context during the movie — transitions from one ‘event’ to another — we used the event boundaries as defined by (Ben-Yakov & Henson, 2018). As part of their study, a sample of 16 participants were asked to watch the movie and to indicate whenever they felt like one event ended and another began. The authors used these subjective ratings to define 19 likely ‘boundaries’ in the movie. For the purpose of the current analyses, we used these times to create event transition windows, defined as the boundary TR +/-2 TRs to capture any gradual as well as abrupt changes in context around the boundary, additionally shifted forward by 2 TRs (∼5s) to account for the hemodynamic lag.

### 2.3. MRI data acquisition

The CamCAN MRI data were collected using a Siemens 3T TIM Trio scanner, with a 32 channel head coil, at the MRC Cognition and Brain Sciences Unit, Cambridge, UK. Functional data during movie watching were acquired with a multi-echo T2* EPI sequence over 193 volumes [32 axial slices, 3.7mm thick, 0.74mm gap, TR = 2470ms, TE = [9.4, 21.2, 33, 45, 57] ms, flip angle = 78 degrees, FOV =192 × 192mm, voxel size = 3 × 3 x 4.44mm]. T1 images were acquired with a 3D MPRAGE sequence [TR = 2250ms, TE = 2.99ms, TI = 900ms, flip angle = 9 degrees, FOV = 256 x 240 x 192m, 1mm isotropic voxels, GRAPPA acceleration factor = 2]. Fieldmap scans were additionally collected [TR = 400ms, TE = 5.19ms/7.65ms, 1 Magnitude and 1 Phase volume, 32 axial slices, 3.7mm thick, 0.74mm gap, flip angle = 60 degrees, FOV = 192 × 192mm, voxel size = 3 × 3 × 4.44mm].

### 2.4. FMRI data processing

The description of MRI data processing below was taken from the custom language generated by fMRIPrep (Esteban et al., 2018), which has been released under the CC0 licence and is recommended for use in publications.

MRI data was preprocessed using fMRIPrep 1.5.2; https://fmriprep.org/en/stable/; RRID:SCR_016216), which is based on Nipype 1.3.1 (Gorgolewski et al., 2011); https://nipype.readthedocs.io/en/latest/; RRID:SCR_002502). Many internal operations of fMRIPrep use Nilearn 0.5.2 (https://nilearn.github.io/; RRID:SCR_001362). The T1-weighted (T1w) image was corrected for intensity non-uniformity with N4BiasFieldCorrection, distributed with ANTs 2.2.0 (http://stnava.github.io/ANTs/, RRID:SCR_004757), and was then skull-stripped with a Nipype implementation of the antsBrainExtraction.sh workflow (from ANTs), using OASIS30ANTs as target template. Brain tissue segmentation of cerebrospinal fluid (CSF), white-matter (WM) and gray-matter (GM) was performed on the brain-extracted T1w using fast (FSL 5.0.9; https://fsl.fmrib.ox.ac.uk/fsl/fslwiki; RRID:SCR_002823). Volume-based spatial normalization to the MNI152NLin6Asym template was performed through nonlinear registration with antsRegistration (ANTs 2.2.0), using brain-extracted versions of both T1w reference and the T1w template.

For the functional data, a reference volume and its skull-stripped version were generated using a custom methodology of fMRIPrep. A deformation field was estimated based on the field map that was co-registered to the BOLD reference, which was used to correct for susceptibility distortions. Head-motion parameters with respect to the BOLD reference (transformation matrices, and six corresponding rotation and translation parameters) were estimated before any spatiotemporal filtering using mcflirt (FSL 5.0.9). BOLD runs were slice-time corrected using 3dTshift from AFNI (https://afni.nimh.nih.gov/; RRID:SCR_005927). The BOLD time-series (including slice-timing correction) were resampled onto their original, native space by applying a single, composite transform (using antsApplyTransforms) to correct for head-motion and susceptibility distortions. A T2* map was estimated from the preprocessed BOLD by fitting to a monoexponential signal decay model with log-linear regression. For each voxel, the maximal number of echoes with reliable signal in that voxel were used to fit the model. The T2* map was used to optimally combine preprocessed BOLD across echoes. The combined time series was carried forward as the preprocessed BOLD, and the T2* map was also retained as the BOLD reference. The BOLD reference was then co-registered to the T1w reference using flirt (FSL 5.0.9) with the boundary-based registration cost-function and nine degrees of freedom. The BOLD time-series were resampled to the MNI template with 2mm voxel resolution.

FMRIPrep calculates several confounding time-series based on the preprocessed BOLD. Framewise displacement (FD) and DVARS were calculated for each functional run, both using their implementations in Nipype (following the definitions by Power et al., 2014). Three global signals were extracted within the CSF, the WM, and the whole-brain masks. Additionally, a set of physiological regressors were extracted to allow for component-based noise correction using the CompCor method (Behzadi et al., 2007). Principal components were estimated after high-pass filtering the preprocessed BOLD time-series (using a discrete cosine filter with 128s cut-off) for the two CompCor variants: temporal (tCompCor) and anatomical (aCompCor). tCompCor components are calculated from the top 5% variable voxels within a mask covering the subcortical regions. aCompCor components are calculated within the intersection of the aforementioned mask and the union of CSF and WM masks calculated in T1w space, after projection to the native space of each functional run.

After preprocessing with fMRIPrep, the confounds were inspected to determine if data met the criteria for inclusion. Subjects were excluded if more than 20% of time points exceeded a FD of 0.3mm and/or if the mean FD exceeded 0.2mm. After careful visual inspection of the data, subjects were additionally excluded if notable artifacts were present or preprocessing had failed. The CONN v.18.b toolbox (Whitfield-Gabrieli & Nieto-Castanon, 2012); https://web.conn-toolbox.org/; RRID:SCR_009550) was used to denoise the BOLD time-series with nuisance regression prior to analyses. For each subject, confound time-series included in the model were the six head motion parameters and their temporal derivatives, the first six aCompCor components from a combined WM and CSF mask, and FD. Additional spike regressors were included for any time points that exceeded a FD of 0.6mm and/or a standardized DVARS of 2. The mean number of spikes identified across subjects was 1.61 (max=13) out of 193 time points. After regression of motion confounds, BOLD data were band-pass filtered with a high-pass filter of 0.008 Hz and a low-pass filter of 0.1Hz. BOLD data were kept unsmoothed for extracting the mean time-series from regions of interest (ROIs), but were smoothed with a 6mm FWHM kernel for whole-brain seed-to-voxel connectivity analyses.

### 2.5. Regions of interest

We used a combination of functional and anatomical atlases to accurately delineate our PMN ROIs. Posterior medial regions within two previously characterized default subnetworks were selected from a cortical atlas (Schaefer et al., 2018) — labeled ‘Default A’ and ‘Default C’ — which reflect regions that are broadly associated with constructive and episodic processes (Andrews-Hanna et al., 2010, 2014; DiNicola et al., 2020). We additionally included the posterior hippocampus (body and tail) from a probabilistic parcellation (Ritchey, Montchal, et al., 2015) due to its well-known role in episodic memory, sensitivity to event boundaries (Reagh et al., 2020), and connectivity to cortical PMN regions (Libby et al., 2012). Next, we used “episodic”-related activity, as defined by a Neurosynth meta-analysis (Yarkoni et al., 2011), to search for a single functional peak within each regional mask, except for medial parietal cortex regions (covering precuneus and posterior cingulate) where two peaks separated by at least 10 voxels were identified. To create each ROI, 100 contiguous (adjoining faces) episodic-sensitive voxels (2 x 2 x 2mm) were selected that expanded out from a peak, constrained by the regional mask. This resulted in 8 equal-sized clusters (Figure 1a): posterior hippocampus (pHipp), parahippocampal cortex (PHC), retrosplenial cortex (RSC), precuneus (Prec), posterior cingulate cortex (PCC), posterior angular gyrus (pAG), anterior angular gyrus (aAG), and medial prefrontal cortex (MPFC).

**Fig 1.**
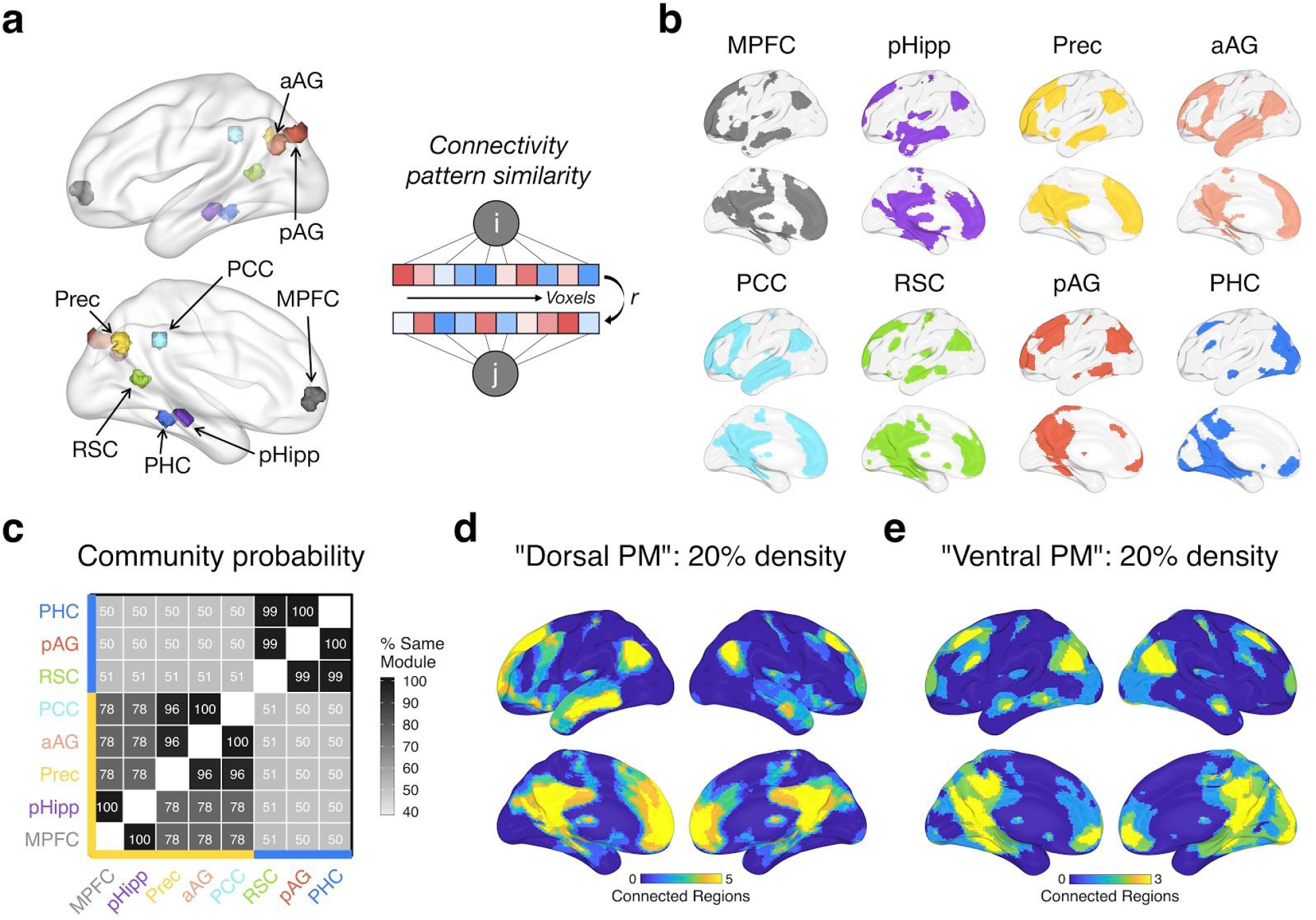
Subsystems of the PMN derived from seed-to-voxel connectivity patterns. a) Left: PMN ROIs, showing each 100-voxel cluster. aAG = anterior angular gyrus, pAG = posterior angular gyrus, PCC = posterior cingulate cortex, MPFC = medial prefrontal cortex, pHipp = posterior hippocampus, PHC = parahippocampal cortex, RSC = retrosplenial cortex, Prec = precuneus. Right: Connectivity pattern similarity approach — subsystems were characterized based on the similarity (Pearson’s r) of whole-brain seed-to-voxel connectivity patterns for each pair of ROIs (i, j). b) The top 20% of group-averaged connections (binarized) between each ROI (seed) and every voxel across the brain. c) Using connectivity density thresholds between 10% and 30%, Louvain community detection was run on the similarity of group-averaged voxel connectivity patterns with gamma values between 0.75 and 1.25. The matrix shows the percentage of the time each pair of ROIs were assigned to the same module across all density and gamma iterations. ROIs are grouped with hierarchical clustering for visualization, which reveals 2 subsystems: The yellow axis line shows regions within a “Dorsal PM” subsystem and the blue axis line shows regions within a “Ventral PM” subsystem. d&e) The overlap in binarized connections shown in (b) for Dorsal PM (d) regions (MPFC, pHipp, Prec, aAG, PCC) and for Ventral PM (e) regions (RSC, pAG, PHC). Warmer colors show a higher number of regions within connections to a voxel. Data is plotted from Group 1 (discovery sample) only — Group 2 (replication sample) results look almost identical, revealing the same subnetworks (Supplementary Analyses S.1).

### 2.6. Statistical analyses

All code is available through our github repository: http://www.thememolab.org/paper-camcan-pmn/. Data were analyzed using MATLAB, R v3.5.1, and RStudio v1.0.143. Brain images were generated with BrainNet Viewer (Xia et al., 2013), and all other plots were generated with ggplot2 within the R tidyverse (https://www.tidyverse.org/). The analyses outlined below tested the functional connectivity of the PMN during movie watching, and how its connections relate to events in the movie. First, we used patterns of whole-brain connectivity to distinguish two PMN subsystems, assessing the strength of intranetwork functional connectivity within and between these subsystems. Next, we tested how the PMN subsystems are similarly or differentially modulated by the content of the movie, including events transitions. Finally, in an exploratory analysis, we investigated how move-related PMN functional connectivity relates to individual differences in episodic memory.

#### 2.6.1. PMN subsystems from functional connectivity patterns

For each region of the PMN, seed-to-voxel connectivity values were calculated as the Pearson’s correlation between the mean ROI time-series (averaged over voxels in the unsmoothed data) and each voxel’s time-series (from smoothed data) during the movie-watching functional scan. This resulted in a whole brain connectivity map per ROI and subject. Subject-level connectivity maps were averaged (after Fisher’s z transformation), per ROI, to produce group-level maps. Louvain community detection (Blondel et al., 2008), from the Network Toolbox (Christensen, 2018), was used to identify likely PMN subnetworks based on the similarity of group-averaged ROI voxel connectivity patterns, applied iteratively over different connection density thresholds and gamma values. Specifically, each ROI connectivity map was binarized according to density thresholds of 10-30%, by 5% increments, such that the top X% of voxel connections to the seed were marked as 1, with connections below that threshold marked as 0. The correlation between the binarized, group-averaged connectivity maps was calculated between every pair of ROIs, and this similarity matrix was run through the Louvain algorithm with gamma values between 0.75 and 1.25, in .01 increments, with gamma influencing how coarse or fine the returned community structure will be. Each iteration of community detection, with a unique density and gamma combination, assigned the PMN ROIs to mutually exclusive groups, and the probability of every pair of ROIs being assigned to the same group was calculated across all iterations. Therefore, our ROI groupings reflect the most robust community structure when considering different levels of voxel connectivity strength. Hierarchical clustering was applied to the matrix of shared probabilities to determine the most appropriate groupings of ROIs into subsystems.

Intranetwork functional connectivity over the entire duration of the movie was calculated as the pairwise Pearson correlations among the 8 ROI time-series for every subject. Subject-level correlation matrices were Fisher-z transformed prior to group-averaging. Mean functional connectivity within and between the PMN subsystems was compared with paired-sample t-tests to validate the distinction between subsystems at the intranetwork level. Supplementary analyses were performed to probe the importance of individual PMN regions in mediating intranetwork connectivity (see Supplementary Analyses).

#### 2.6.2. Movie-related dynamic functional connectivity

Whereas the previous functional analyses were based on time-averaged connectivity across the movie, remaining analyses targeted the relevance of *time-varying* PMN connectivity in relation to the movie input. First, we considered the influence of event transitions in the movie. For each pair of ROIs, we entered their standardized time-series as well as their product (coactivation), reflecting the edge time-series (Faskowitz et al., 2019; van Oort et al., 2018), as predictors in a logistic regression model with event type (transition:1 vs. within-event:0) as the dependent variable [*event ∼ ROIi + ROIj + ROIi*ROIj*]. Event transition time points were defined as a boundary TR +/-2 TRs to capture any gradual as well as abrupt changes in context around the boundary, additionally shifted forward by 2 TRs (∼5s) to account for the hemodynamic lag. The beta coefficient of the interaction term reflects the change in ROI coactivation at event transitions, controlling for the activity time-series of each ROI. These beta values were averaged within and between PMN subsystems to test the sensitivity of PMN connections to event transitions.

Second, we considered whether fluctuations in PMN connectivity are tied to the movie stimulus (including but not limited to event transitions), which is shared across subjects, by comparing the intersubject similarity of time-varying connectivity. Within each subject and for every time point *t*, we first calculated the Spearman correlation between every pair of ROI time-series over a window of 25 TRs (∼62s) centered on *t*, resulting in a vector of time-varying connectivity. This window size is consistent with prior work characterizing time-varying correlations during movie-watching (Di & Biswal, 2020; Jang et al., 2017; Simony et al., 2016), with the aim of smoothing noise that accompanies the raw edge time-series or correlations over very short windows, while preserving temporal resolution, before comparing time-varying connectivity between subjects (Di & Biswal, 2020). To test if PMN connectivity fluctuations are tied to the movie, we next calculated the intersubject Pearson correlation of time-varying connectivity vectors (Di & Biswal, 2020), averaged within each subsystem, using a leave-one-subject-out approach (Nastase et al., 2019): For each subject, we calculated the correlation between their time-varying connectivity and the average time-varying connectivity of all remaining subjects, resulting in one intersubject correlation value per subject. If connectivity fluctuations are meaningfully related to the movie content (shared across subjects), then the average intersubject correlation should be non-zero. We also compared the intersubject correlations of the same subsystem to intersubject correlations across different subsystems to test if movie-related dynamics of PMN subsystems are distinct from one another. All r values were Fisher-z transformed prior to averaging.

#### 2.6.3. Relationship between PMN connectivity and episodic memory

In a final exploratory analysis, we tested how PMN connectivity relates to episodic memory on an independent task. Within our full sample of CamCAN subjects, exactly half (N=68) had also completed a separate item-scene memory task, where neutral objects were paired with a negative, neutral, or positive background scene. In a memory test, participants recalled the objects and their associated scene context, verbally describing the details of the scene (see Shafto et al., 2014) for a detailed task description). One subject was excluded from analyses due to a high number of response errors (> 50% of trials). We used the number of neutral trials for which the scene context was recalled in detail as a measure of episodic memory for each subject. As a control measure, we used priming of objects in the neutral condition, as indexed by corrected recognition of previously studied degraded objects.

We first considered the influence of time-averaged connectivity, testing if there was a correlation between mean PMN subsystem functional connectivity and episodic memory across subjects. Second, we considered the influence of time-varying connectivity by using intersubject representational similarity analysis (IS-RSA), as outlined by Finn et al., 2020). A subject x subject representational dissimilarity matrix (RDM) was calculated for behavioral scores, and another was calculated for brain data. The brain RDM was defined as the intersubject dissimilarity (1 -Pearson correlation) of time-varying connectivity (or activity, see Supplementary Analyses) for each of the PMN subsystems. Two behavioral RDMs were tested — a nearest neighbor model, reflecting the Euclidean distance between the memory scores of every pair of subjects [*abs(i-j)*], and an “Anna K” model (as coined by Finn et al., 2020), where high performing subjects are assumed to be similar, with increasing variability among lower performing subjects [*max score -min(i,j)*]. Finn and colleagues found that the Anna K model, using behavioral data from a working memory task, provided the best fit to their brain data. Spearman correlation was then calculated between every behavioral RDM and brain RDM. To determine the significance of the behavior-brain correlations, 10,000 permutations were run, wherein the subject labels for the brain RDM were shuffled for every permutation. The *p*-value for the behavior-brain comparison was calculated as the proportion of permutation correlations that were greater than the true correlation. Bonferroni-corrected *p*-values were also calculated, correcting for the total number of tests (6: 2 behavioral models x 3 time-varying connectivity measures).

## 3. Results

### 3.1. PMN subsystems from functional connectivity patterns

First, we tested whether regions in the PMN were dissociable based on their patterns of connections with the rest of the brain during movie watching. To do so, we examined seed-to-voxel connectivity across the whole brain, using the similarity of whole-brain connectivity patterns between ROIs (Figure 1a) to group them into subsystems with Louvain community detection. Comparing the similarity of seed-to-voxel connectivity patterns across PMN ROIs revealed a high degree of overlap in the strongest connections. As expected, voxels within the broader default network, including medial prefrontal cortex, medial and lateral parietal cortex, and lateral temporal cortex, were consistently within the top 20% of functional connections to PMN regions (Figure 1b), supporting their common grouping within a coherent network. However, two distinct PMN subsystems were identified (Figure 1c). We tested the replicability of these results in a second group of subjects, which revealed an identical allocation of ROIs to PMN subsystems (see Supplementary Analyses S.1).

Across multiple connection thresholds and Louvain gamma values (see Methods 2.6.1), PHC, pAG, and RSC were grouped into the same module almost 100% of the time, hereafter referred to as the “Ventral posterior medial (PM)” subsystem (Figure 1e). These regions shared a module assignment with other ROIs approximately 50% of the time. On the other hand, MPFC, pHipp, Prec, aAG, and PCC were assigned to the same module at least 78% of the time, hereafter referred to as the “Dorsal PM” subsystem (Figure 1d). Due to this high overlap and limited number of ROIs, we grouped all 5 of these regions together, but it is interesting to note that pHipp and MPFC were particularly similar in their voxel connectivity patterns, being grouped into the same module 100% of the time. The same was true for Prec, aAG, and PCC, suggesting that a finer-grained parcellation may be possible.

Next, we examined functional connectivity within the set of PMN ROIs. Like the seed-to-voxel analyses, Pearson’s correlations between the ROI time-series confirmed a high-degree of interconnectedness (Figure 2a). Despite significant connectivity between virtually all pairs of ROIs, evidence for the subsystems identified on the basis of whole-brain connectivity was supported (Figure 2b) — on average, connections within both the Ventral and Dorsal PM subsystems were stronger than connections between them (ts(67) > 4.32, *p*s < .001; replicated in Group 2: ts(67) > 4.06, *p*s < .001).

**Fig 2.**
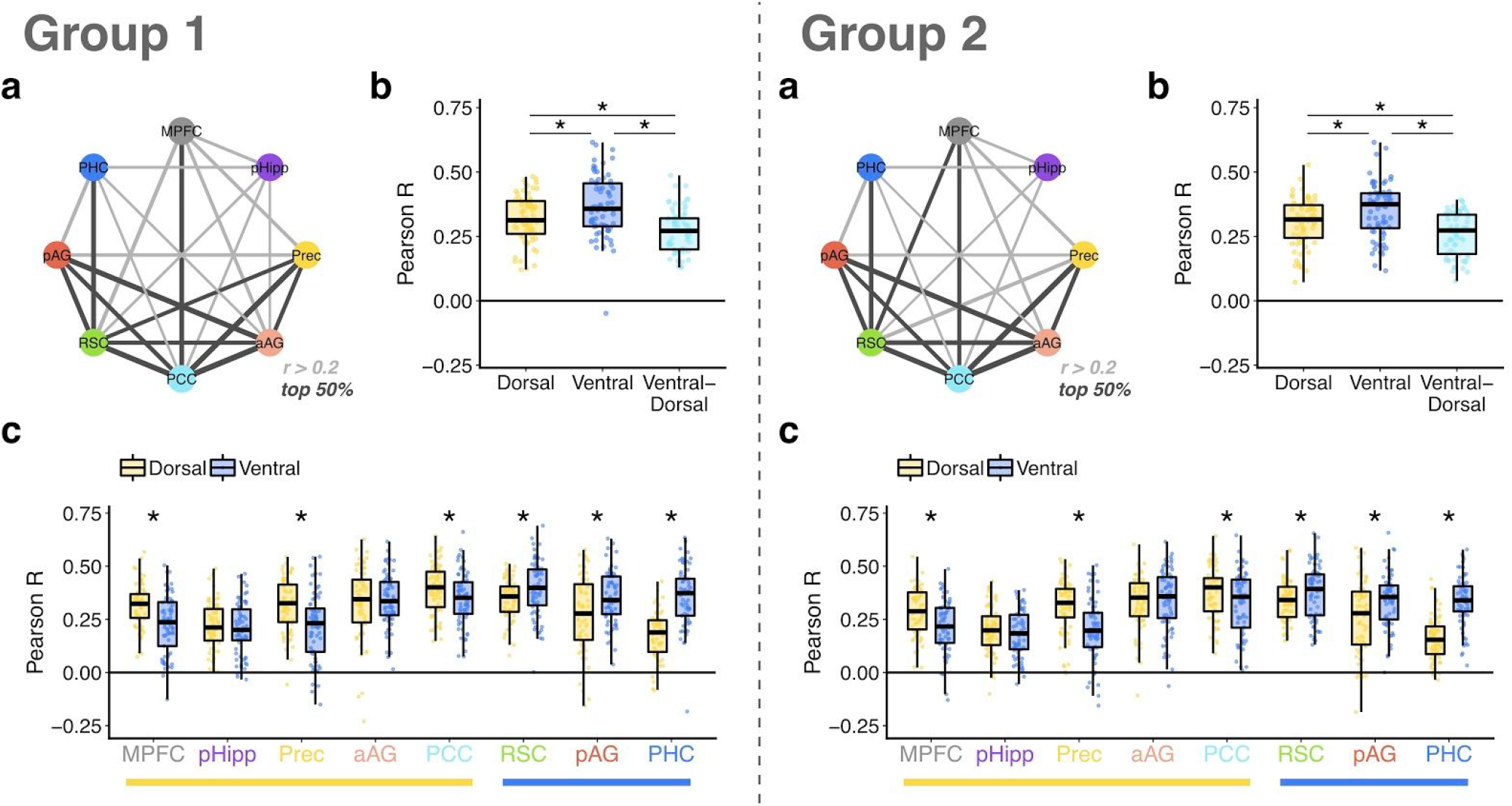
Intranetwork PMN functional connectivity. Left = Group 1 (discovery sample) results; right = Group 2 (replication sample) results. a) Graph showing intranetwork PMN connections: light gray edges indicates mean r > .2, dark edges indicate the top 50% of those connections. A mean r value of 0.2 was deemed to be a conservative threshold for visualizing a meaningful edge as it reflects the critical value for a significant correlation (at *α* = .005) between our ROI time-series with 193 TRs. b) Distribution of mean functional connectivity within and between PMN subsystems (each point indicates a subject). c) Distribution of mean functional connectivity for every ROI to the two PMN subsystems. The 5 ROIs to the left, underlined in yellow, form the Dorsal PM subsystem. The 3 ROIs to the right, underlined in blue, form the Ventral PM subsystem. * indicates a significant difference between subsystem connectivity strength at *p* < .05.

We additionally calculated the average strength of functional connectivity between each PMN region and each of the subsystems (Figure 2c). All 3 ROIs within the Ventral PM network (RSC, pAG, PHC) showed significantly stronger functional connectivity to each other than to regions of the Dorsal PM network (ts(67) > 3.02, *p*s < .004; replicated in Group 2: ts(67) > 2.21, *p*s < .031). Within the Dorsal PM network, MPFC, Prec, and PCC showed stronger communication with other Dorsal PM regions than with the Ventral PM subsystem (ts(67) > 2.55, *p*s < .013; replicated in Group 2: ts(67) > 2.28, *p*s < .026); however, pHipp and aAG did not show preferential connectivity to one subsystem over another (ts(67) < 1.02 *p*s > .31; replicated in Group 2: ts(67) < 0.98, *p*s > .32). This likely reflects the integration of pHipp with PHC and the strong communication between aAG and pAG, despite the whole-brain connectivity patterns of pHipp and aAG being more aligned with other Dorsal PM regions. In contrast, PHC and Precuneus showed the strongest, opposite differentiation in functional connectivity between the PMN subsystems. Overall, RSC and PCC had the strongest functional connectivity within the PMN, which is supported by their large, yet distinct, mediating effect on PMN connections: In a supplementary analysis, we found that RSC mediated communication between PHC and the Dorsal PM subsystem, whereas PCC mediated communication among Dorsal PM regions (see Supplementary Analyses S.2).

### 3.2. Movie-related dynamic functional connectivity

The prior results show the presence of two PMN functional subsystems and reveal key nodes that mediate their communication through analyses of time-*averaged* functional connectivity over the entire movie. Here, we sought to validate the functional significance of these subsystems in terms of their time-*varying* dynamics. First, we tested how connectivity within and between the subsystems changed at event transitions in the movie. In line with prior analyses of event boundaries in the CamCAN dataset (Ben-Yakov & Henson, 2018; Reagh et al., 2020), we confirmed a general increase in activity across all PMN ROIs at event transitions (see Supplementary Results S.3). Therefore, we asked how connectivity is modulated by event transitions over and above these activity changes.

We found a striking and replicable dissociation between the subsystems in the sensitivity of connectivity to event transitions. Whereas coactivation within the Dorsal PM subsystem did not change as a function of event transitions (Group 1: *t*(67) = -0.94, *p* = .35; Group 2: *t*(67) = 0.07, *p* = .94), there was a strong increase in coactivation within the Ventral PM subsystem (Group 1: *t*(67) = 8.46, *p* < .001; Group 2: *t*(67) = 9.26, *p* < .001), and between Ventral and Dorsal PM regions (Group 1: *t*(67) = 5.66, *p* < .001; Group 2: *t*(67) = 7.51, *p* < .001) (see Figure 3a&b). In both groups, the change in functional connectivity of the Ventral PM subsystem at event transitions was significantly greater than changes within the Dorsal PM subsystem (Group 1: *t*(67) = 7.57, *p* < .001; Group 2: *t*(67) = 7.55, *p* < .001) and between Ventral and Dorsal PM regions (Group 1: *t*(67) = 6.10, *p* < .001; Group 2: *t*(67) = 4.91, *p* < .001). We additionally verified that the event transition window length (defined as a 5-TR window centered on each HRF-adjusted boundary) did not drive our effects: repeating the analyses using shorter 3-TR windows did not change the selective modulation of Ventral PM connectivity.

**Fig 3.**
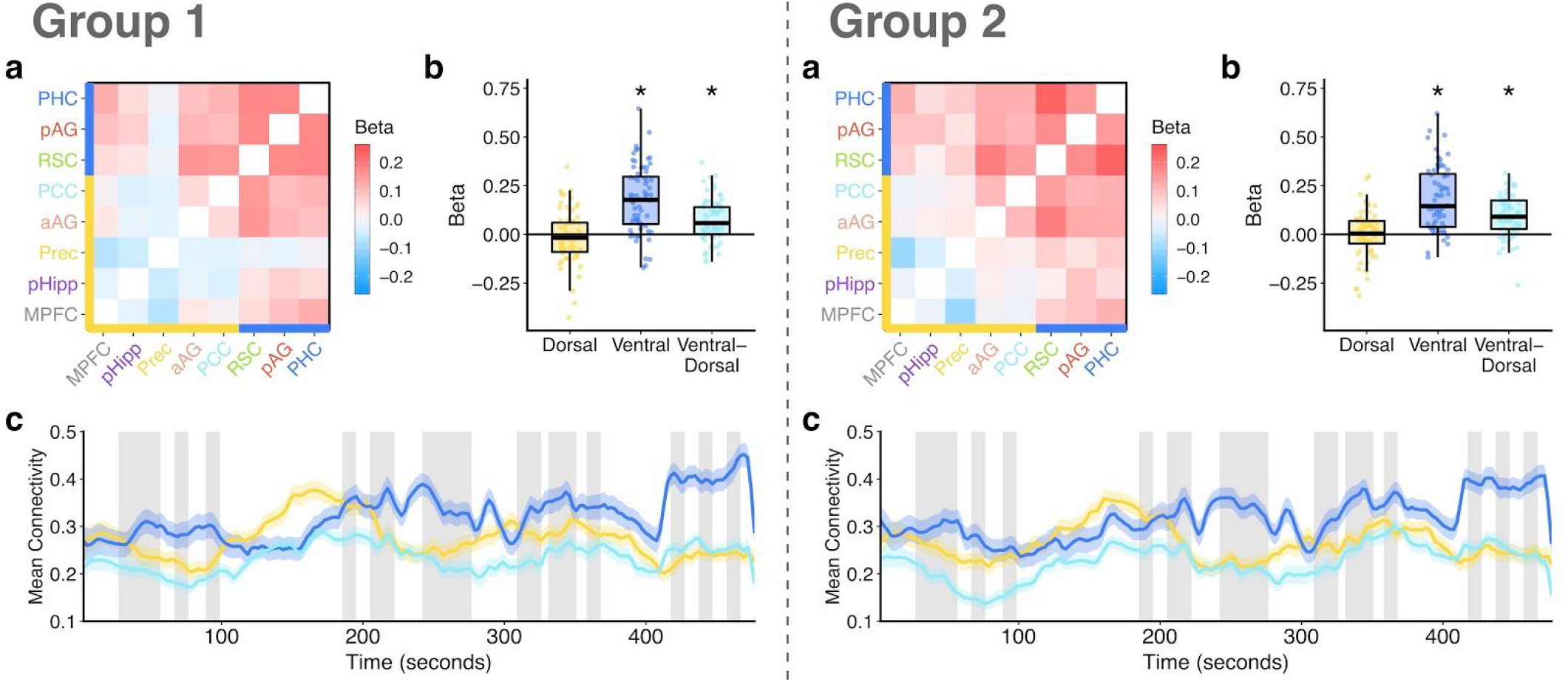
Change in PMN connectivity at event transitions. Left = Group 1 (discovery sample) results; right = Group 2 (replication sample) results. a) The group-averaged beta values, reflecting the change in connectivity between each pair of ROIs at event transitions relative to within events. Warmer colors reflect an increase in connectivity at transitions. Yellow axis line shows Dorsal PM regions; blue axis line shows Ventral PM regions. b) The distribution of mean subsystem changes in connectivity at event transitions. Each point indicates a subject, * indicates a change significantly greater than zero at *p* < .05. c) Group-averaged time-varying connectivity is plotted for connections within and between the PMN subsystems. Line = mean across subjects, ribbon = standard error of the mean. Gray windows indicate event transitions within the movie (shifted by 2TRs to account for the HRF). Note that some transition windows are immediately adjacent to one another, producing wider windows. Time-varying connectivity is calculated using a 25-TR sliding window, centered on each TR.

To probe movie-related connectivity dynamics within and between the PMN subsystems beyond their specific relation to event transitions, we additionally calculated time-varying connectivity using a sliding-window (Figure 3c). We then compared the similarity of time-varying connectivity across subjects to determine if PMN subsystem connectivity was related to the movie content (shared across subjects) and if connectivity of the two PMN subsystems exhibited similar or distinct fluctuations. Interestingly, visual inspection of the data shows that the only point during the movie where group-averaged connectivity among Dorsal PM regions exceeded that among Ventral PM regions was during a prolonged event, without any transitions, from approximately 100s to 182s. Intersubject correlations revealed that connectivity fluctuations of the whole PMN appear to be tied to the movie content, as evidenced by significant intersubject similarity of time-varying connectivity for both the Ventral PM subsystem (Group 1: mean Z = 0.23, SE = 0.04, t(67) = 6.03, *p* < .001; Group 2: mean Z = 0.20, SE = 0.04, t(67) = 5.45, *p* < .001) and the Dorsal PM subsystem (Group 1: mean Z = 0.30, SE = 0.05, t(67) = 6.74, *p* < .001; Group 2: mean Z = 0.25, SE = 0.04, t(67) = 5.57, *p* < .001). In contrast, there was not a positive intersubject relationship between the time-varying connectivity of the Ventral PM subsystem and the time-varying connectivity of the Dorsal PM subsystem (Group 1: mean Z = -0.09, SE = 0.03, t(67) = -3.04, *p* = .003; Group 2: mean Z = -0.02, SE = 0.02, t(67) = -1.01, *p* = .32). Therefore, connectivity fluctuations within PMN subsystems appear tied to the movie input as shown by consistency across subjects, rather than reflecting only idiosyncratic patterns, but these fluctuations differed between the subsystems, suggesting that they were related to different aspects of the movie.

Visual inspection of the data shows a notable increase in time-varying connectivity of the Ventral PM subsystem just after 400 seconds (see Figure 3c). Interestingly, this phase of the movie is centered around the main event — an accidental gunshot. To verify that this period did not drive any of our results, we repeated all analyses only including data up to TR 162/193 (400 seconds). None of the findings changed when using this shortened version of the data.

### 3.3. Relationship between PMN connectivity and episodic memory

The prior analyses demonstrated functional differences between PMN subsystem connectivity patterns and their relation to the movie events. But what is the significance of PMN connectivity for memory-related behavior? To gain some preliminary insight into this question, we conducted exploratory analyses testing if functional connectivity related to individual differences in episodic memory performance on an independent task (Figure 4). In this task, participants were tested on their memory for the scene context associated with studied objects. Episodic memory was defined as the number of trials for which participants could describe the scene context in detail.

**Fig 4.**
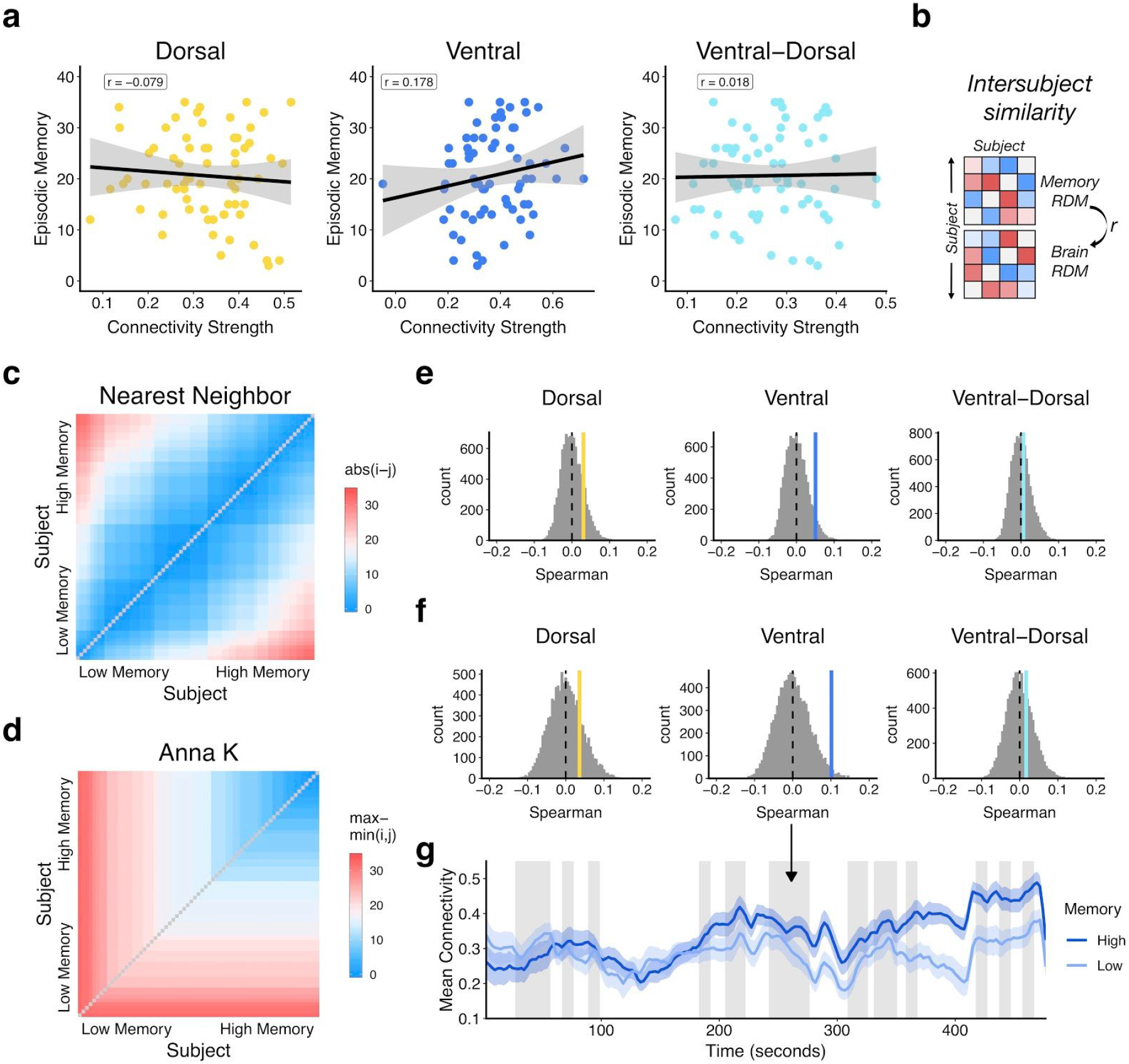
Relationship between PMN connectivity and episodic memory. a) Correlations between time-averaged functional connectivity and episodic memory. b) Intersubject representational similarity calculation, wherein a memory representational dissimilarity matrix (RDM) is correlated with a brain RDM. c) The memory RDM using a nearest neighbor model, where subjects with similar memory scores are assumed to have similar brain dynamics. d) The memory RDM using an “Anna K” model (Finn et al., 2020), where subjects with high memory are assumed to have similar brain dynamics, with low memory subjects assumed to be more variable. e&f) Null distributions of Spearman correlations between memory and brain RDMs over 10,000 permutations, shuffling subject labels in the brain RDM. The solid, colored lines, indicate the true correlation between the behavior and brain RDMs for each PMN subsystem. e) Permutations of the nearest neighbor model, f) Permutations of the “Anna K” model. g) Mean time-varying connectivity of the Ventral PM subsystem for high and low episodic memory subjects, defined by a median split (ribbon = standard error of the mean). Gray windows = event transitions for visualization.

There was no significant relationship between time-averaged PMN subsystem connectivity strength during the movie and episodic scores across subjects (Figure 4a; Dorsal: r = -.079, *p* = .52; Ventral: r = .178, *p* = .15; Ventral-to-Dorsal: r = .018, *p* = .88). Next, we used intersubject representational similarity analysis (Figure 4b; Finn et al., 2020) to test the relationship between time-varying connectivity and episodic memory. We tested two models — nearest neighbor (Figure 4c), where subjects with similar behavior are assumed to have similar connectivity dynamics, and an “Anna K” model (Figure 4d), where high performing subjects are assumed to have similar connectivity, with low performing subjects being more variable (see Finn et al., 2020). A nearest neighbor model (Figure 4e) revealed no significant relationship between time-varying PMN connectivity and episodic memory (Dorsal: r = .030, *p* = .15; Ventral: r = .051, *p* = .053, Bonferroni-corrected *p* = .32; Ventral-to-Dorsal: r = .007, *p* = .37). However, the Anna K model (Figure 4f) suggested a positive, selective relationship between the intersubject similarity of episodic memory and Ventral PM time-varying connectivity (r = .101, *p* = .017) although we highlight that this effect did not survive correction across all 6 models (Bonferroni-corrected *p* = .10). This correlation was not present for time-varying Dorsal (r = .036, *p* = .20) or Ventral-to-Dorsal PM connections (r = .017, *p* = .30). Control analyses showed no significant correspondence between time-varying PMN processes and memory for activity or a measure of object priming from the same task (see Supplementary Analyses S.4). We also verified that there was no relationship between subjects’ episodic memory performance and their movement (mean FD) during the movie watching scan (r = -.04).

To visualize any potential correspondence between Ventral PM subsystem connectivity and episodic memory, the mean time-varying connectivity was calculated for subjects with high (N=35) vs. low (N=32) memory scores, using a median split (Figure 4g). This revealed distinct changes in Ventral PM connectivity over the movie between memory groups: post-hoc tests for a linear trend in time-varying connectivity showed that subjects who had high episodic memory scores significantly increased their Ventral PM connectivity over time (mean Z = 0.326, t(34) = 5.31, *p* < .001), whereas subjects with lower scores did not (mean Z = 0.002, t(31) = 0.04, *p* = .97), which was significantly different between groups (t(64.6) = 3.67, *p* < .001). Therefore, not only do PMN subsystems show meaningfully distinct patterns of functional connectivity during movie watching, those dynamics, particularly of Ventral regions, may have implications for individuals’ episodic memory.

## 4. Discussion

The PMN is a structurally and functionally interconnected system that specializes in the construction of episodic representations. Prior work has provided valuable insight into the functional properties and representations of the PMN as a whole, but research is only just starting to map the organizational structure of the PMN, which is important for understanding the multidimensional nature of episodic thought (Ritchey & Cooper, 2020). To this end, we sought to tease apart functional pathways and subsystems within the PMN, testing how they relate to changes in event context and individual differences in episodic memory. First, we found that the PMN can be parcellated into two functional subsystems: a Ventral PM system comprising PHC, RSC, and posterior AG, and a Dorsal PM system comprising PCC, Prec, anterior AG, MPFC, and posterior Hipp. Second, we showed that functional connectivity with PHC and Prec differentiated the subsystems, whereas RSC and PCC were strongly connected throughout the PMN. Third, we found that although connectivity of both the Dorsal PM and Ventral PM subsystems tracked the movie content, showing time-varying similarity across subjects, there was a selective increase in functional connectivity of Ventral PM regions at event transitions. Finally, time-varying connectivity of the Ventral PM subsystem appeared to relate to individual differences in episodic memory.

The partition of the PMN into two separable functional subsystems during movie watching aligns with prior work considering fractionation of the broader default network during rest. Early approaches identified an MTL default subsystem, characterized by strong connectivity to PHC, that included RSC and posterior AG, and a Dorsal Medial subsystem, comprising dorsal MPFC and lateral temporal cortex, with these two subsystems converging on a Core subsystem of dorsal medial parietal cortex and MPFC (Andrews-Hanna et al., 2010). Recent work in individuals, however, has divided the default network into two interdigitated subsystems (Braga & Buckner, 2017; DiNicola et al., 2020) that retain notable overlap with groupings of PMN regions identified here. In particular, default network ‘A’ is characterized by strong functional connectivity to PHC, whereas default network ‘B’ includes regions within anterior lateral parietal cortex, PCC, and MPFC that appear similar to our Dorsal PM network. However, these parcellations have not included the hippocampus and other subcortical structures, and the individual-specific analyses do not afford a direct comparison with the current results or with group-level functional dissociations in the literature. More recently, the default network has been segregated beyond two systems, including a separate Parietal Network of middle AG, medial parietal cortex, and anterior MPFC, and a Ventromedial Network of hippocampus and ventral MPFC (Barnett et al., 2020; Gordon et al., 2020). We also observed some evidence for a functional divide within the Dorsal PM system, with reliable clustering of hippocampus and MPFC based on similar patterns of whole-brain connectivity. Interestingly, these prior studies and our results suggest that the hippocampus may be more strongly aligned with the network organization of MPFC despite strong functional communication with PHC. Overall, the subsystems we revealed within the PMN during movie-watching appear to align with those that have been identified within the default network during rest using a whole-brain approach. An outstanding question is whether movie-watching or other episodic paradigms might be best suited to studying PMN organization because they directly modulate the network.

Beyond the overarching network organization, analyses of intranetwork functional connectivity validated the separation of PMN subsystems and revealed the connectivity patterns of individual PMN regions. The hippocampus consistently exhibited the lowest functional connectivity suggesting that, whereas PHC and connections of the Dorsal PM system — most notably MPFC — converge on the hippocampus, it may not drive communication among cortical PMN regions. In support, prior work suggests that temporal integration of narrative information during movie watching in Dorsal PMN regions may not depend on interactions with the hippocampus (Chen et al., 2016; Zuo et al., 2020). The hippocampus has been characterized as a gateway between the PMN and an anterior temporal (AT) network that processes item and emotional information (Ranganath & Ritchey, 2012; Ritchey, Libby, et al., 2015). Therefore, while it may not be central to situation models supported by cortical PMN regions, the hippocampus may be important for connecting the PMN with other brain networks. In contrast, RSC and PCC were the most strongly functionally connected regions within the PMN, but our analyses highlighted distinct roles: RSC mediated communication between PHC and the Dorsal PM subsystem, whereas PCC mediated communication among Dorsal PM regions. The dominance of both RSC and PCC in the network supports prior work that has highlighted these areas as connectivity “hubs”. PCC is often regarded as an integrative hub of the default network (Andrews-Hanna et al., 2010; Buckner et al., 2008), as revealed with partial correlation analyses (Fransson & Marrelec, 2008), that can regulate information flow between default regions (Wang et al., 2019). Complementing the current findings, RSC in turn has been shown to mediate connectivity between the MTL and dorsal default network regions (Kaboodvand et al., 2018), and is thought to serve as a key area of transformation between MTL and dorsal parietal spatial codes (Bicanski & Burgess, 2018).

In addition to understanding the functional structure of the PMN, we sought to demonstrate the functional relevance of subsystems for event processing. Prior work has shown increased activity throughout the PMN at event boundaries (Ben-Yakov & Henson, 2018; Reagh et al., 2020), as replicated here. However, we observed a selective increase in connectivity of the Ventral PM subsystem, not among Dorsal PM regions, at event transitions. This dissociation highlights both a distinct finding from connectivity patterns that is not observed with regional activity alone, and a dominant role of Ventral PM communication in integrating events within the PMN. Interestingly though, time-varying connectivity of *both* the Ventral and Dorsal PM subsystems was ‘synced’ to the movie stimulus, as evidenced by intersubject correlations, but they were unrelated to one another. This suggests that the subsystems were related to different features of the movie, and thus the event-transition dissociation between subsystems is not reflective of any overall difference in the sensitivity of connectivity fluctuations to movie content. The movie shown to participants was 8 minutes in total, with some events lasting only a few seconds. A fine-grained sensitivity to event structure among Ventral PM regions supports findings of a temporal event hierarchy in the PMN: RSC and PHC process the most high-resolution events in contrast to more slowly evolving context models in Dorsal PM regions (Baldassano et al., 2017; Keidel et al., 2017), as perhaps indicated by the increase in Dorsal PM connectivity during the longest event in our analysis. Moreover, within the default network, connectivity between the medial temporal lobe and RSC, specifically, increases during episodic tasks relative to rest (Bellana et al., 2017).

We suggest that, at local event boundaries, Ventral PM regions communicate with Dorsal PM regions to integrate the event with a sustained and more abstract contextual framework. This explanation is in line with evidence of intra-PMN dissociations in the reinstatement of event context, which is persistent in AG, Prec, and PCC and more transient in PHC and RSC (Jonker et al., 2018). An outstanding question, therefore, is whether within-Dorsal PM connectivity increases at boundaries characterized by less frequent thematic shifts that could not be explicitly modeled with the current task. Relatedly, it is unclear whether Ventral PM connectivity is particularly sensitive to boundaries that are characterized by shifts in visuo-spatial content or whether it reflects a content-general process that would be sensitive to other kinds of context shifts, such as semantic narrative. Providing some support for the latter, a prior study suggests that RSC and PHC are sensitive to changes in narrative context when visuo-spatial context is maintained (Keidel et al., 2017). Finally, a surprising finding was the lack of increase in hippocampal connectivity at event transitions, particularly given the dominant role of hippocampal event boundary signals in supporting memory (Cohen et al., 2015; Cooper & Ritchey, 2020; Reagh et al., 2020). It is possible, however, that the posterior hippocampus creates and separates event-specific representations (Chanales et al., 2017; Schlichting et al., 2015) in contrast to the integrated event structure (embedding specific events within an ongoing situational model) supported by cortical PMN communication (Aly et al., 2018).

Mirroring the modulation of PMN connectivity by movie event transitions, an exploratory analysis showed some evidence for a selective relationship between Ventral PM connectivity and episodic memory on an independent task. Subjects who had detailed recollection of scene context showed more similar patterns of time-varying Ventral PM connectivity during movie watching, which reflected an increase in connectivity over time. Ventral areas of the PMN, particularly PHC and RSC, are strongly related to the processing of spatial contextual information in memory and imagination in contrast to the preferential representation of people in posterior cingulate cortex and anterior angular gyrus (Peer et al., 2015; Robin et al., 2018; Silson et al., 2019). PHC, RSC, and posterior AG have also been defined as a functionally connected system supporting scene memory (Baldassano et al., 2016; Steel et al., 2020). Moreover, functional connectivity of Ventral PM regions, specifically, has been previously related to episodic memory: One study found that PHC-and RSC-mediated resting state connectivity of the hippocampus and AG was related to TMS-enhanced spatial memory precision (Tambini et al., 2018). In another study, there was a selective relationship between MTL-RSC resting state connectivity and episodic memory that was not present for other default network connections (Kaboodvand et al., 2018). In contrast to this prior research, we did not find a relationship between time-averaged connectivity and individual differences in memory. Rather, we provide preliminary evidence that Ventral PM connectivity *dynamics* may be relevant for individual differences in episodic memory, although we highlight that the main effect did not survive Bonferroni correction. Moreover, the cause of the relative dissimilarity in time-varying connectivity for subjects with low episodic memory is unclear, possibly reflecting differences in general attention and processes not specific to memory. While we speculate that increased Ventral PM connectivity over time in subjects with high episodic memory could indicate an enhanced ability to bind spatial-contextual information, future research will be required to test this hypothesis and, importantly, replicate the current results.

In conclusion, we revealed distinct functional subsystems of the PMN, whose pathways dynamically tracked movie content. Communication of the Ventral PM subsystem was selectively modulated by event transitions during movie-watching and may relate to individual differences in episodic memory. Beyond these specific findings, our analyses point to both the utility and challenges of integrating large-scale networks with questions about specific cognitive operations, which are often studied in a region-centric manner. Understanding the subnetwork organization of brain networks, and mapping that organization to specific task-related factors, may be the key to understanding the functional relevance of large-scale networks associated with high-level cognitive processes (Cabeza et al., 2018; Cabeza & Moscovitch, 2013; Ritchey & Cooper, 2020). Overall, our findings illustrate PMN functional organization, and highlight the significance of functional diversity of the PMN for event cognition.

## Acknowledgements

Data collection and sharing for this project was provided by the Cambridge Centre for Ageing and Neuroscience (CamCAN). The authors also thank Aya Ben-Yakov for making the movie event boundary data available. CamCAN funding was provided by the UK Biotechnology and Biological Sciences Research Council (grant number BB/H008217/1), together with support from the UK Medical Research Council and University of Cambridge, UK. S.W.D. was supported by NIH grant K01AG053539 and M.R. was supported by NIH grant R00MH103401.

## Supplementary Analyses

### S.1 Replication of PMN subsystems

The seed-to-voxel connectivity patterns were highly similar between Group 1 (shown in Fig.1, main paper) and the replication Group 2 (Fig.S1). The replication sample resulted in identical allocation of PM ROIs to two subsystems, also suggesting the potential for a more refined structure of the Dorsal PM subsystem with an alliance between pHipp and MPFC and a robust grouping of Prec-aAG-PCC.

**Fig S1.**
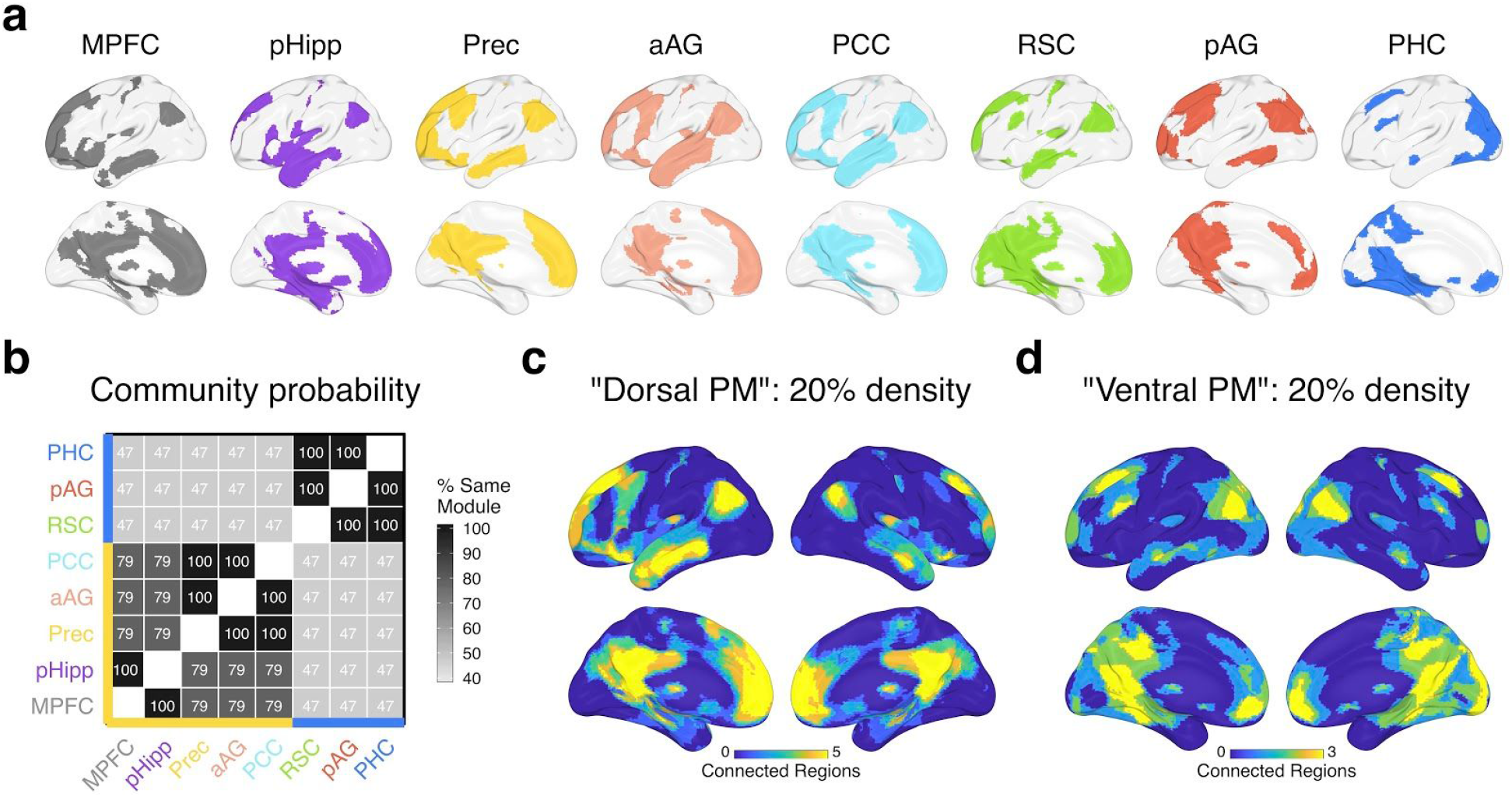
Subsystems of the PMN from seed-to-voxel connectivity patterns: Replication (Group 2) results. a) The top 20% of group-averaged connections (binarized) between each seed and every voxel across the brain. b) The percentage of the time of each pair of ROIs were assigned to the same module across all density and Louvain gamma values. The yellow axis line shows regions within a “Dorsal PM” subsystem and the blue axis line shows regions within a “Ventral PM” subsystem. c&d) The overlap in binarized connections shown in (a) for Dorsal PM (c) regions (MPFC, pHipp, Prec, aAG, PCC) and “Ventral PM” (d) regions (RSC, pAG, PHC).

### S.2 Mediating influence of PMN ROIs on intranetwork connectivity

In order to visualize the mediating influence of each ROI on intranetwork functional connectivity, we controlled for each ROI prior to calculating functional connectivity among remaining PMN regions. Specifically, for every ROI *m*, the partial correlation between the residuals of all other pairs of ROIs was calculated after removing the variance explained by *m* (see Kaboodvand et al., 2018). The remaining network thus reflects the connectivity structure after statistically removing one region. The partial correlation values (*c’*) were then compared to the original, bivariate correlations (*c*) to calculate the proportion of connectivity that is mediated by *m* [*Pm = 1 -(c’/c)*]. At the subject-level, the overall influence of *m* on PMN connectivity is calculated by contrasting the mean of *c* and *c’* across all network edges. At the group-level, the mean values of *c* and *c’* across subjects were contrasted per edge to illustrate the specific network pathways that *m* mediates. For this edge-specific analysis, edges were considered only if their group-level bivariate correlation value (*c*) exceeded .2. Prior to averaging, both *c* and *c’* were Fisher-z transformed. For both of the aforementioned subject-level and group-level analyses, we excluded any potential suppressing effects (where *c’* > *c, i*.*e. Pm < 0*) by setting *Pm* to 0 in such instances, and also excluded the influence of mean negative values of *c’ (i*.*e. Pm > 1)* by setting Pm to 1 in these cases. Therefore, proportion mediated was constrained to the range 0-1.

**Fig S2.**
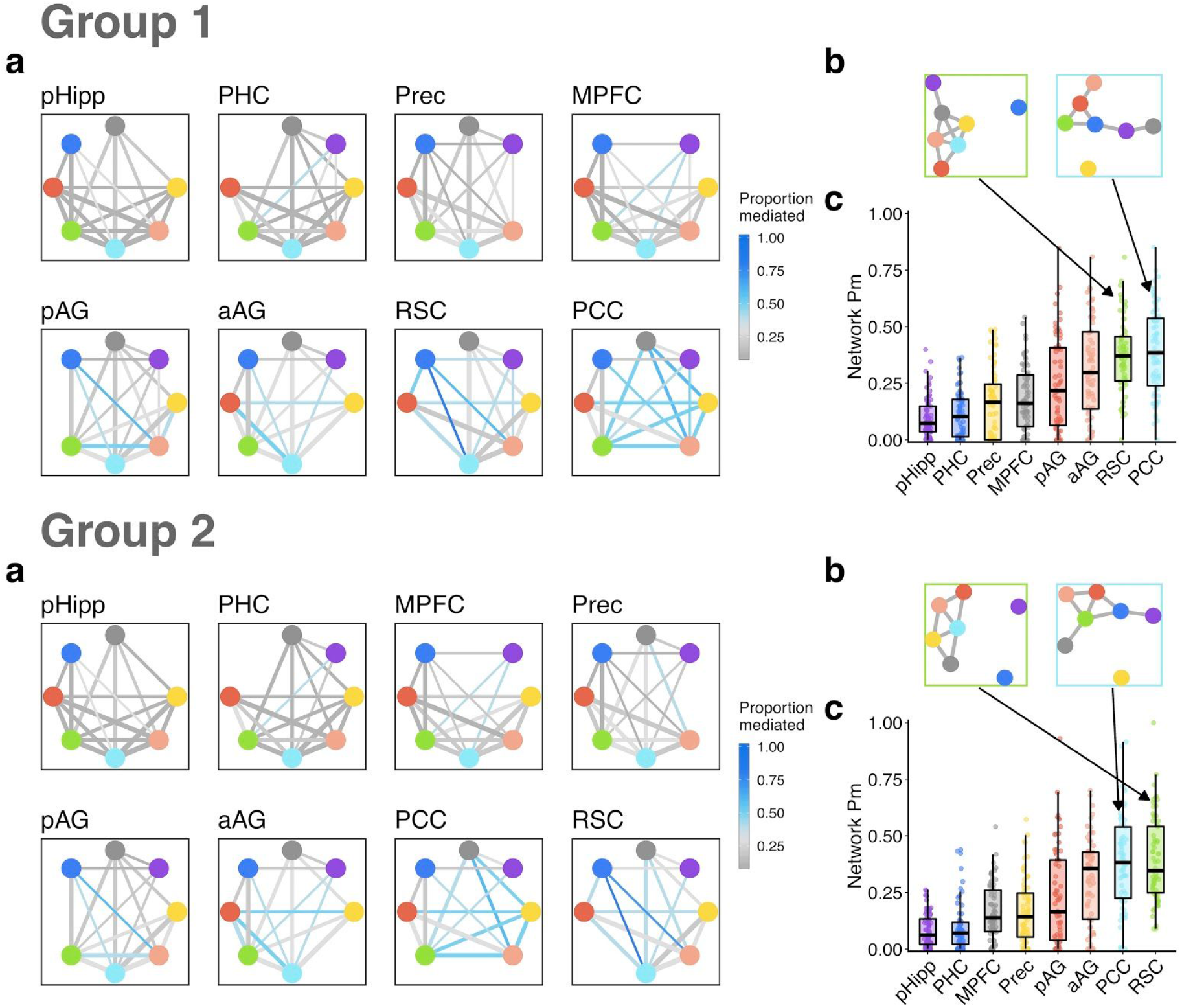
Mediating influence of PMN ROIs. Top = Group 1 (discovery sample) results; bottom = Group 2 (replication sample) results. a) The effect of removing the variance explained by each ROI on PMN edges. Plotted edges reflect all group-averaged bivariate correlations (r) > .2, with width reflecting the original, bivariate edge strength. Edge color indicates the proportion of the edge that is mediated by the ROI labelled in the top left of each panel (removed from the accompanying graph), such that blue colors indicate a larger reduction in an edge’s strength relative to gray colors. Proportion mediated (Pm) values for edges are calculated based on group-averaged data. b) The effect of RSC (left) and PCC (right) on connectivity between remaining PMN regions, showing edges (mean r) that are > .2 after statistical removal of the ROI. c) For each subject (point), the proportion of mean connectivity across the rest of the network that is mediated by the ROI along the x axis. Panels in (a) are sorted by total network Pm as shown in (c).

These results revealed a number of interesting patterns. First, pHipp, PHC, Prec, and MPFC have a small mediating influence on the rest of the PMN (top row, Figure S2a), replicated in Group 2. Therefore, even though these regions communicate with one another and other PMN regions, few intranetwork connections are mediated by their activity. There were two notable exceptions — PHC influenced connectivity of pHipp to RSC, and MPFC also partially mediated connectivity of pHipp to RSC and PCC. Second, both anterior and posterior AG exerted a moderate influence on PMN connectivity, but with specific roles: pAG mediated communication between the Ventral PM system (PHC and RSC) and aAG. In turn, aAG mediated connectivity between pAG and dorsal medial parietal regions — Prec and PCC. Finally, RSC and PCC exerted the strongest influence on PMN connectivity, but with complementary roles: RSC mediated communication between the Ventral and Dorsal PM subsystems, with a particularly large influence on PHC pathways, but left communication between Dorsal PM regions largely intact. In contrast, PCC has almost no effect on connectivity among Ventral PM regions but substantially mediated communication among Dorsal PM regions, with the exception of pHipp-MPFC, as well as integration of the Dorsal PM subsystem with RSC.

### S.3 Sensitivity of PMN activity to event transitions

The current analyses use the event boundaries identified by (Ben-Yakov & Henson, 2018) in their original analysis of the CamCAN movie-watching data. With a larger sample than that included here (due to a higher upper age limit) and different analysis methods (including larger ROIs from the Harvard-Oxford atlas), the authors showed increased activity of the hippocampus, PHC, and posterior medial cortex (including Prec and PCC/RSC) in response to event boundaries. To validate this finding with our methods and ROIs, the mean change in activity from within-event to event-transition time points was calculated for each subject from their z-scored time series, per ROI. Our results replicate those of Ben-Yakov & Henson as well as (Reagh et al., 2020) in showing robust increases in activity at event boundaries across the PMN (Figure S3).

**Fig S3.**
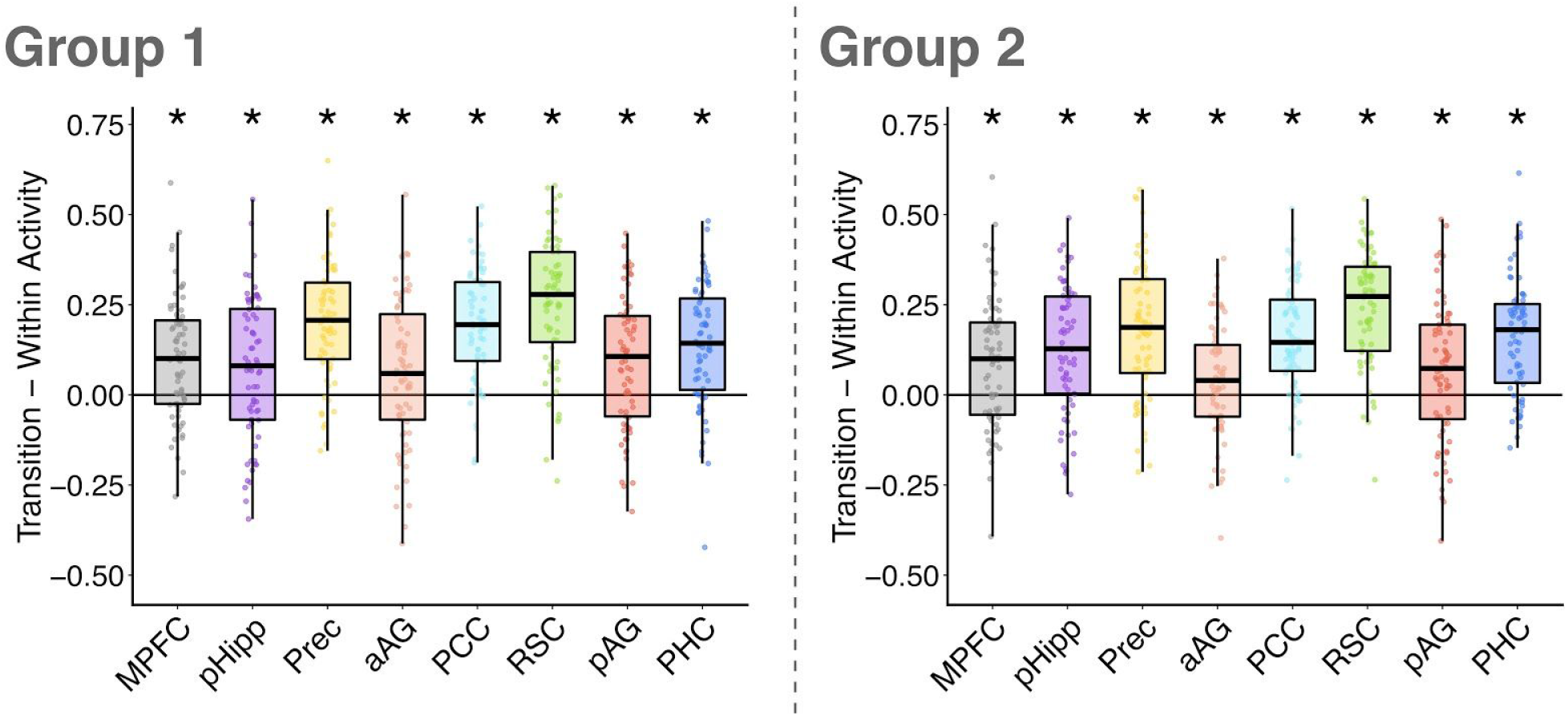
The change in activity between within-event time points and event transitions. ROI time series were first z-scored within each subject and mean activity was contrasted between phases. Points indicate individual subjects, * indicates *p* < .05, FDR-corrected across all ROIs. Results reveal a consistent increase in activity across the PM network at event transitions, that is replicated in group 2.

### S.4 Intersubject representational similarity control analyses

To test the specificity of the intersubject representational similarity analysis results, we ran two control analyses — one to test if there was relationship between time-varying PMN subsystem *activity* and episodic memory, and the other to test if intersubject similarity of time-varying PMN connectivity related to an independent memory measure from the same task, object priming. Using the same method as for analyses of time-varying connectivity, intersubject similarity of activity was defined as the correlation of mean PMN subsystem activity time-series for every pair of subjects. There was no significant relationship between the intersubject similarity of PMN activity and episodic memory using both the nearest neighbor model (Dorsal: r = .028, *p* = .25; Ventral: r = -.004, *p* = .52) and the Anna K model (Dorsal: r = .078, *p* = .14; Ventral: r = .071, *p* = .19). In testing the relationship between the intersubject similarity of time-varying PMN connectivity and object priming, we also found no significant relationships using both the nearest neighbor model (Dorsal: r = .037, *p* = .12; Ventral: r = -.020, *p* = .71; Ventral-to-Dorsal: r = .019, *p* = .24) and the Anna K model (Dorsal: r = .053, *p* = .11; Ventral: r = -.045, *p* = .85; Ventral-to-Dorsal: r = -.058, *p* = .97). Therefore, a correspondence between time-varying Ventral PMN processes and individual differences in memory appears to be specific to connectivity and to episodic performance within this task.

